# Selective dissociation of vacant 70S ribosomes by RsfS promotes bacterial adaptation to nutrient starvation

**DOI:** 10.1101/2024.03.23.586405

**Authors:** Hong Zhang, Rainer Nikolay, Liang Jiang, Yariv Brotman, Gong Zhang, Bo Qin

## Abstract

Bacteria rapidly remodel protein synthesis to survive nutrient limitation, yet how inactive ribosomes are selectively regulated remains incompletely understood. Ribosome silencing factor S (RsfS) is the only known bacterial factor that selectively dissociates vacant 70S ribosomes, but its physiological function in bacterial adaptation has remained elusive. Here, we demonstrate that RsfS promotes bacterial survival during nutrient starvation by selectively dismantling non-translating vacant 70S ribosomes while sparing translating ribosomes. Biochemical analyses show that RsfS efficiently dissociates vacant 70S ribosomes but has little effect on programmed ribosomes in the pre-translocation or post-translocation states, thereby preserving ongoing protein synthesis. Ribosome profiling further reveals that loss of RsfS results in a global reduction in translational efficiency and widespread translational reprogramming during starvation. Mechanistically, dissociation of vacant ribosomes by RsfS suppresses futile EF-G-dependent GTP hydrolysis and prevents unnecessary energy expenditure associated with inactive ribosome cycling. Consistent with this mechanism, RsfS enhances bacterial adaptation to nutrient deprivation without compromising active translation. Together, our findings identify RsfS as a unique regulator of bacterial ribosome homeostasis that conserves cellular energy through selective dissociation of vacant 70S ribosomes, revealing a previously unrecognized strategy by which bacteria optimize translation under nutrient-limited conditions.

## Introduction

For optimal bacterial growth, active translation is imperative to synthesize proteins. Translational regulation is crucial for prompt responses to environmental shifts, such as nutrient shortage, oxidative stress, accumulation of toxic metabolites, and quorum sensing, given the faster regulatory dynamics at the translational level compared to the transcriptional level (Mukherjee, Raghavan, & Chatterji, 1998; Njenga, Boele, Öztürk, & Koch, 2023; Smits, Kuipers, & Veening, 2006; Waters & Bassler, 2005; Zhao, Qin, Nikolay, Spahn, & Zhang, 2019; Zhong et al., 2015). The ribosome serves as the cellular machinery for protein translation, necessitating various factors to ensure both accuracy and efficiency (Wilson & Nierhaus, 2007). Nevertheless, the mechanisms and biological significance of growth-phase-dependent downregulation of protein synthesis by noncanonical factors remain enigmatic. For instance, ribosomes isolated from stationary-phase cells exhibit lower protein synthesis activity compared to those from logarithmic-phase cells (Bommer et al., 1996). Despite the identification of numerous factors that inhibit translation (Franken et al., 2017; Izutsu et al., 2001; Prossliner, Skovbo, Sørensen, & Gerdes, 2018; Ueta et al., 2013), this phenomenon still remains poorly understood.

The stringent response is an extensively studied mechanism that enables bacteria to cope with nutrient deprivation. In this response, RelA and SpoT in *E. coli* regulate the synthesis of components of the translational apparatus to inhibit the *de novo* ribosome synthesis under amino acid starvation *via* the alarmone (p)ppGpp (Arenz et al., 2016; Christensen, Mikkelsen, Pedersen, & Gerdes, 2001; Kudrin et al., 2018; Loveland et al., 2016; Wendrich, Blaha, Wilson, Marahiel, & Nierhaus, 2002). However, the number of existing ribosomes remains unaltered during stringent response. It is known that the growth rate of bacteria strongly correlates with the number of ribosomes present (Ehrenberg, Bremer, & Dennis, 2013; Gausing, 1977), increasing linearly as the number of ribosomes in the cell rises from 7,000 to 70,000 (H Bremer & Dennis, 1987). Therefore, the down-regulation of the number of ribosomes per cell reduces cellular resource consumption under starvation conditions, promoting bacterial survival. Additionally, polysomal fractions are severely reduced or even disappear, and single 70S ribosomes become the dominant fraction *in vivo* (Dresden & Hoagland, 1967a, 1967b). Moreover, it has been demonstrated that the translation elongation rate is maintained at a surprisingly high level (>50% of that in rich medium) even during extremely slow growth in poor medium, indicating that the overall translational apparatus is downregulated by limiting the pool of active ribosomes in the cell. This regulation of translation is physiologically important for the effective synthesis of proteins when nutrients are depleted (Dai et al., 2016). A number of ribosome-associated factors participate in this regulatory process; some induce an inactive, hibernating state of the ribosome in the form of 70S monomers (RaiA) or 100S dimers (RMF and HPF homologs) (Prossliner et al., 2018). Recent studies have further expanded the repertoire of bacterial ribosome hibernation factors, highlighting the diversity of ribosome preservation strategies under stress. However, these factors primarily stabilize inactive ribosomes, whereas mechanisms for selectively eliminating vacant 70S ribosomes remain largely unknown (Helena-Bueno et al., 2024). RsfS has been identified as the sole factor that affects the equilibrium between 70S ribosomes and their subunits (Häuser et al., 2012).

RsfS and its homologs are present in almost all bacteria and organelles but not in Archaea, and are involved in translational regulation (Häuser et al., 2012). Structural and interaction studies of RsfS and its mitochondrial ortholog MALSU1 reveal their binding affinity to the large ribosomal subunit and specific interaction with the ribosomal protein uL14 (Butland et al., 2005; Gavin et al., 2006; Häuser et al., 2012; Khusainov et al., 2020; Li et al., 2015; Rorbach, Gammage, & Minczuk, 2012; Wanschers et al., 2012). Although the deletion of *rsfS* in *E. coli* is not lethal (Baba et al., 2006), it leads to significantly impaired viability during the stationary growth phase, along with a growth arrest for approximately 20 hours during the transition from nutrient-rich to nutrient-poor medium, resulting in these bacteria being out-competed when co-cultured with WT bacteria (Häuser et al., 2012). Moreover, loss of MALSU1 induces malfunctions in the mitochondrial respiratory system (Rorbach et al., 2012). Recent work in the intracellular pathogen *Legionella pneumophila* has further underscored the physiological importance of RsfS, demonstrating that it functions coordinately with other hibernation factors to support starvation survival, host cell infection, and antibiotic tolerance (Schmid et al., 2026). Given its specific binding site and inhibitory effect of translation *in vivo* and *in vitro*, RsfS has been suggested to block translation by preventing the formation of active 70S ribosomes from its subunits (Häuser et al., 2012; Khusainov et al., 2020). However, the detailed mechanism through which RsfS contributes to these regulations and the global molecular consequences remain elusive.

In this study, we show the indispensable role of *rsfS* in mediating strain adaptation to nutrient downshifts, attributed to its facilitation of efficient utilization of cellular resources using ribosome profiling. Remarkably, further biochemical experiments provided evidence that RsfS specifically dissociates vacant 70S ribosomes, while leaving translating ribosomes unaffected. This action effectively alleviates the energy burden associated with idle ribosomes, thereby enabling the enhanced expression of enzymes crucial for biosynthetic pathways and energy production during nutrient deprivation. Deciphering the intricate mechanisms underlying RsfS-mediated adaptation processes provides a comprehensive perspective on the bacterial stress responses.

## Results

### Lack of rsfS causes a severe adaptation challenge during nutrient deprivation

After a nutrient shift from rich (LB) to poor (M9) media, the *E. coli* wild-type (WT) strain exhibited sustained exponential growth. Conversely, the strain lacking *rsfS* ceased growth after 5 hours, entering a 23-hour lag phase before resuming growth (Fig. 1a).

**Figure 1:**
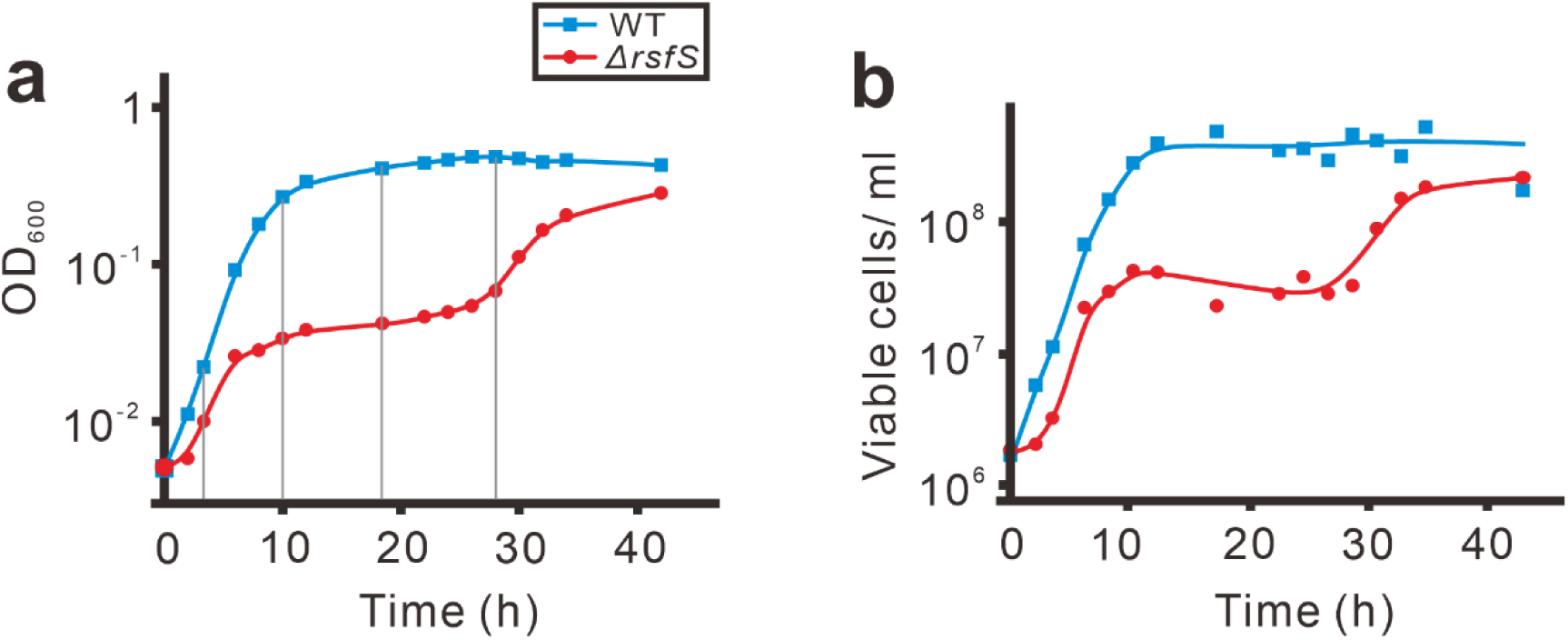
Growth curve and viability test of WT and ΔrsfS strains. WT and *ΔrsfS* strains were cultured overnight in rich medium (LB) and then diluted 1:1000 in poor medium(M9). The cultures were incubated at 37°C for 42 hours. OD_600_ was measured at indicated time points (**a**) and aliquots were spread on LB plates after serial dilutions. After incubation, single colonies were counted and corresponding cell densities were calculated (**b**). The gray lines in (**a**) indicate the time points when aliquots were taken for further analysis. Data are presented as mean values based on n=2 biological replicate.

There are at least three possible explanations for this phenomenon: (i) Lack of RsfS results in comprehensive cell death, with only a small fraction of survivors to generate a new population. (ii) The absence of RsfS induces a strong selection pressure, leading to compensatory mutation(s) that rescue the RsfS phenotype. (iii) This phenomenon represents a significant adaptation challenge, wherein cell growth is arrested and recovery occurs only after overcoming metabolic adaptation hurdles during the transition from rich to poor media.

In the first two cases, the relative viability (viable cells in corresponding OD_600_) is expected to decrease significantly during the growth arrest. In the third case, the relative viability remains stable during the 23-hour period, after which it begins to increase again. Fig. 1b illustrates that number of viable cells reduces slightly during the growth arrest, indicating a significant adaptation challenge for bacterial cells in the absence of RsfS.

### Lack of rsfS results in the accumulation of idle 70S ribosomes in vivo

To elucidate the underlying reasons for growth differences, bacterial cultures of both the WT and *ΔrsfS* strains were isolated at the indicated time points during growth (3h, 10h, 18h, and 28h, Fig. 1a). The cell lysates were then subjected to sucrose gradient ultracentrifugation followed by ribosome profile analysis. In the case of the WT strain, all samples exhibited normal polysome profiles with clear peaks corresponding to 30S, 50S, and 70S ribosomes (Fig. 2a). For clarity, only the profiles of WT samples at 3h and 10h are presented, as the WT strain enters stationary phase after 10h, and the data (10h, 18h, and 28h) are nearly identical. At the fully adapted state (28h), the *ΔrsfS* strain exhibits normal growth, displaying a typical polysome profile similar to that of the WT strain. However, during the lag phase (10h and 18h), the polysome profile reveals nearly undetectable levels of 30S and 50S subunits, with the majority of ribosomes existing in the assembled 70S form (Fig. 2a). This suggests a limited availability of free ribosomal subunits for translation initiation, potentially due to impaired 70S recycling following translation under stress conditions. We observed that the rRNA content of all these samples was nearly identical (Fig. 2b), suggesting that the majority of 70S ribosomes during the lag phase did not dissociate into subunits after completing translation. Instead, they remained in an idle state, meaning they were not actively engaged in translating mRNA but persisted in the 70S form, likely resulting from ribosomal run-off (Zhang et al., 2010).

**Figure 2:**
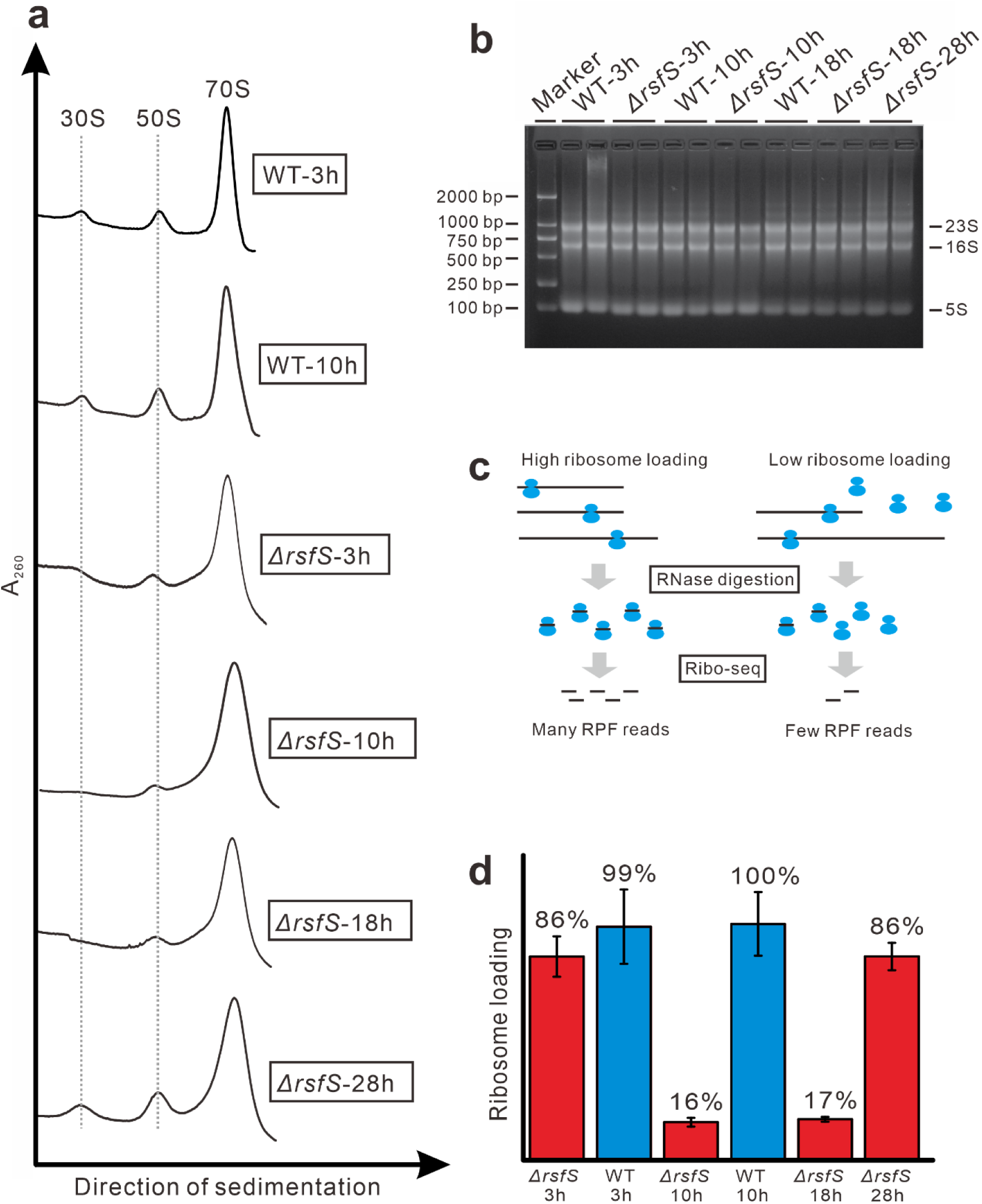
accumulation of idle 70S ribosomes in vivo. (**a**) Ribosome profile in sucrose gradient ultracentrifugation of the samples at representative time points from Fig. 1a. (**b**) Total RNA from the same samples shown in (**a**) resolved by agarose gel electrophoresis. The size marker is a double-stranded DNA marker and ribosomal RNAs are indicated. (**c**) Schematic diagram of ribosome loading (the fraction of 70S ribosomes that are translating mRNAs) measurement. Detailed procedure is illustrated in Fig. S1. RPF: ribosome-protected fragments. (**d**) Results of ribosome loading measurement. The ribosome loading of WT strain at 10h was considered as 100% and all the other samples are normalized accordingly. Data are presented as mean values +/- S.D. based on n=2 technical replicates.

Ribosome profiling (Ribo-seq) experiments were conducted on all these samples to further explore the phenomenon of "idle 70S". We estimated ribosome loading, defined as the fraction of 70S ribosomes engaged in mRNA translation relative to the total 70S fraction, according to the fraction of ribosome-protected fragments (RPF) reads (Fig. 2c). Detailed analysis is provided in the supplementary materials (Fig. S1). We considered the ribosome loading of the WT strain at 10 h as 100% and normalized all the other results accordingly. As expected, during exponential growth (3h), the WT strain exhibited 99% ribosome loading (Fig. 2d), which remained nearly identical at 10 h, suggesting continuous protein synthesis throughout the exponential phase. In contrast, the *ΔrsfS* strain showed 86% loading at 3 h, the initial growth phase, however, its ribosome loading dramatically decreased to 16-17% during the lag phase (10h and 18h), indicating a significant proportion of idle ribosomes. Upon resumption of growth at 28 h, the ribosome loading also returned to 86%, suggesting a robust inverse relationship between ribosome loading and bacterial growth.

### Translational impact of idle 70S ribosomes caused by rsfS absence

Differentially translated genes were screened using Ribo-seq data and edgeR software to explore the global translational impact of *rsfS* (Fig. 3a). The statistically significant overrepresented gene ontology (GO) Biological Process terms of the up-/down-regulated genes at translation level (FDR < 1E-4, listed in Fig. 3b) showed the major physiological changes that occurred when the bacteria were grown in nutrient-depleted conditions. As a positive control, WT bacteria that were grown for 10 hours in poor medium had up-regulated biosynthetic processes including amino acids and organic compounds (Fig. 3b, WT10h vs WT3h). In contrast, in the *ΔrsfS* strain these synthetic processes were severely down-regulated after 10 hours (Fig. 3b, *ΔrsfS*10h vs WT10h). Additionally, in the *ΔrsfS* strain processes relevant to cell motility and localization were also down-regulated reflecting its energy deprivation (Fig. 3b, *ΔrsfS*3h vs WT3h). Direct measurement of the absolute Nucleoside 5’-triphosphates (NTPs) concentration in living cells was hardly to achieve because of all the NTPs can be hydrolyzed to NDPs or NAPs in a couple of seconds in the cell lysate. Therefore, the down-regulated cell motility and localization can serve as an indirect hint for the reduced cellular energy level (Macnab, 1978).

**Figure 3:**
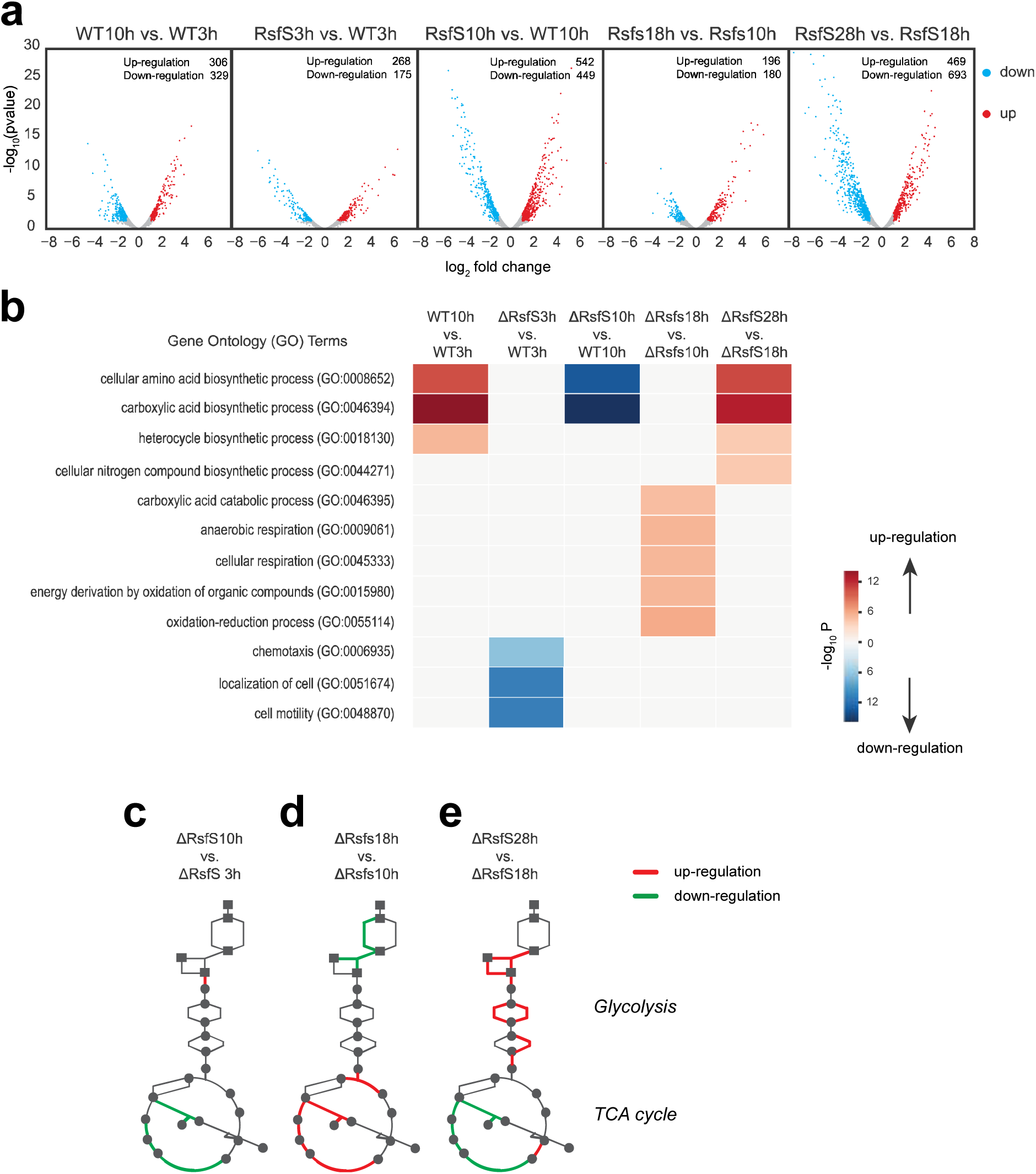
Ribosome profiling demonstrates translatomic impact of RsfS. (**a**) Differential expression analysis of the ribosome profiling sequencing results of the samples. Down-regulated genes were marked as cyan, and up-regulated genes were marked as red. (**b**) Gene Ontology enrichment analysis of the differentially translated genes. Grey blocks indicate non-significant terms (P > 1E-4). All P-values are the FDR values. (**c-e**) The up-/down-regulated genes encoding enzymes in central energy metabolism pathways.

From the results given above, we learned that energy is essential in the process of stress adaptation. Therefore, we next investigated the central energy metabolism pathways (e.g. glycolysis and TCA cycle) for the *ΔrsfS* strain, to understand why the growth and biosynthetic processes were resumed after 28h (Fig. 1a and Fig. 3b, *ΔrsfS*28h vs *ΔrsfS*18h). Although these pathways were not statistically overrepresented in the GO analysis, some of the enzymes in these energy production pathways were still differentially translated. After 18 hours, although the *ΔrsfS* strain still exhibited growth arrest, the TCA cycle was significantly upregulated, as compared to 10 h (Fig. 3c, 3d) which restored the cellular energy for the following recovery (Fig. 3b, *ΔrsfS*18h vs 10h). In sharp contrast, glycolysis in the *ΔrsfS* strain remained unchanged at 10 h, but the TCA cycle was down-regulated (Fig. 3c). This result demonstrated the energy deficiency of the *ΔrsfS* strain at 10 h, and thus unable to synthesize the various enzymes of the biosynthetic processes. The GO analysis showed that the term “energy derivation by oxidation of organic compounds” were significantly overrepresented in the up-regulated genes (FDR= 3.01E-05) (Fig. 3b, *ΔrsfS*18h vs 10h). Taken together, this indicated that the energy production was rescued after 18h. At 28 h, glycolysis genes were largely upregulated compared to 18 h (Fig. 3e). With such energy production, the *ΔrsfS* strain resumed growth, echoed by the eventual up-regulation of various biosynthetic process at 28 h (Fig. 3b). These results suggest that energy deprivation in the *ΔrsfS* strain is the major cause contributing to the deficiency in enzyme synthesis and the growth arrest observed in the initial 10 hours of growth in poor media. In summary, the RsfS factor plays a crucial role in conserving cellular energy under starvation conditions.

### RsfS reduces the cellular translation activity by specifically dissociating idle 70S ribosomes

Translation is one of the most energy- and material-consuming cellular processes. Therefore, it is expected that in nutrient-poor media, translation activity is repressed to conserve energy and resources. This ensures that only essential proteins are expressed, maintaining the translational translocation rate at an appropriate level (Dai et al., 2016). To further investigate the mechanism of RsfS effects on translation, RsfS was added to a transcription-translation lysate assay based on the *E. coli* cell-free system (Roche), in which a plasmid encoding the gene for Renilla luciferase as a reporter was transcribed and translated. As anticipated, the addition of RsfS resulted in an approximately 8-fold reduction in luciferase expression compared to the control, indicating significantly lower translation activity (Fig. 4a). Analytical sucrose gradient ultracentrifugation was performed on the same aliquot from the assay to understand the underlying reason behind this result. The analysis revealed that the addition of RsfS leads to the conversion of 70S ribosomes into their constituent subunits (Fig. 4b). This conversion could be attributed to either the anti-association role of RsfS or its ability to dissociate 70S ribosomes, or perhaps a combination of both mechanisms.

**Figure 4:**
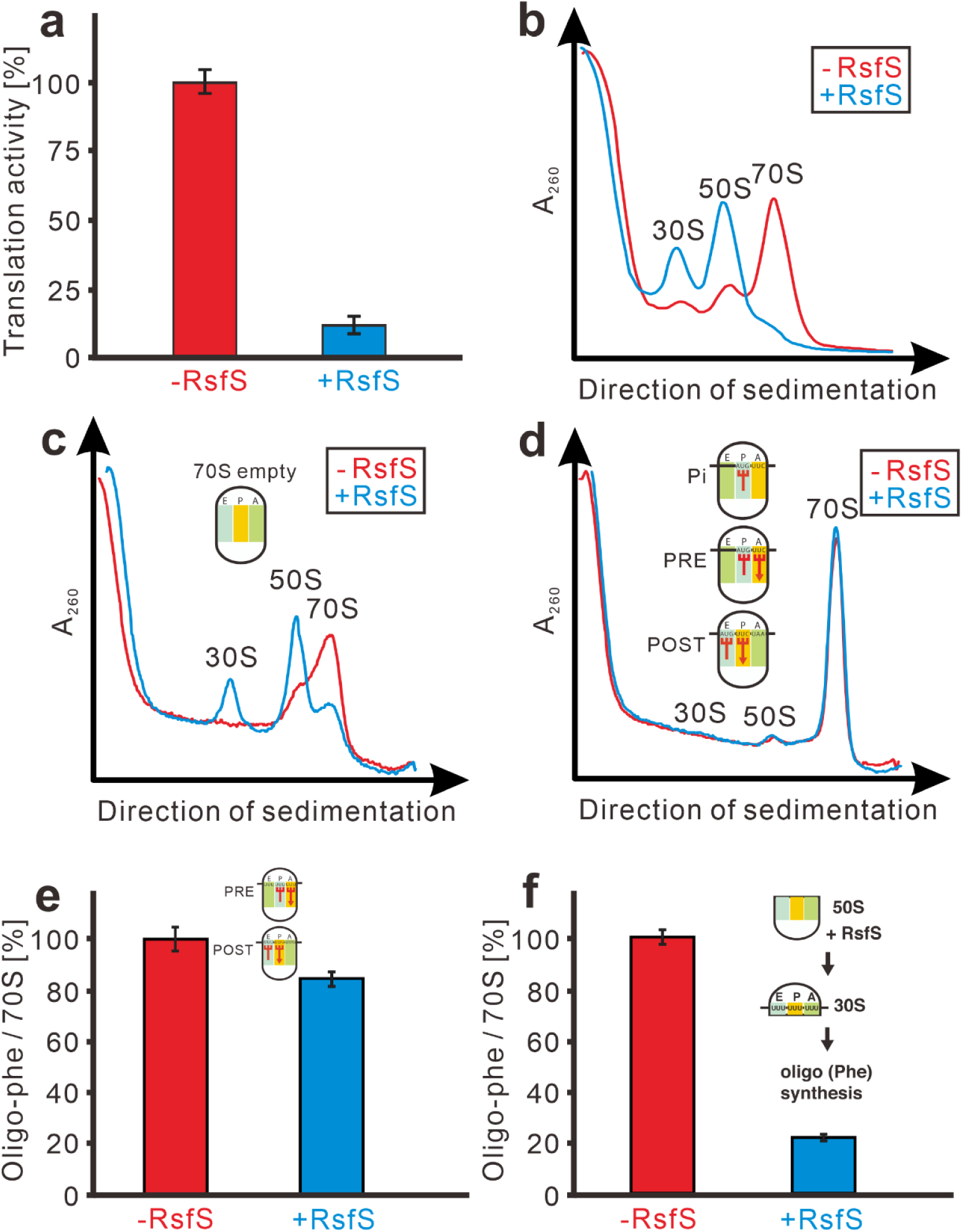
RsfS is an active 70S dissociation factor targeting vacant 70S ribosome. (**a**) Synthesis of Renilla luciferase in a transcription-translation system (RTS). Translation activity was determined by luminescence after 1 h of incubation at 30°C. 10 molar excess of RsfS over ribosomes was used. (**b**) Ribosome profile of the cell lysate translation system used in (**a**). (**c**) RsfS dissociates empty 70S ribosome into ribosomal subunits *in vitro*. RsfS was added in a 10-molar excess over ribosomes. (**d**) RsfS fails to dissociate ribosomes at Pi, PRE and POST states (The functional states of the translating ribosome appearing during the elongation cycle. Pi: i for initiation, programmed ribosomes containing only one tRNA at the P site; PRE: pre-translocational state, characterized by tRNA occupation of both A and P sites; POST: post-translocational state, characterized by tRNA occupation of both P and E sites). The ribosome profiles of these three states are similar. For clarity, we only show here the profile of POST state. RsfS was present in a 10-molar excess. For details see Materials and methods. (**e**) RsfS reduces oligo (Phe) synthesis marginally, when incubated with 70S programmed with mRNA and tRNA. RsfS was added in 10-molar excess over ribosomes. Oligo (Phe) synthesis was determined after an incubation for 10 min at 37°C. (**f**) 50S subunits were preincubated with a 10-fold molar excess of RsfS before 30S, mRNA, [^14^C] Phe-tRNA, GTP and elongation factors were added. The reaction mixture was incubated for 10 min at 37°C, the mixture was analyzed for oligo (Phe) synthesis. Data are presented as mean values +/- S.D. based on n=2 technical replicates.

To confirm the ability of RsfS to dissociate 70S ribosomes, chemically crosslinked 70S ribosomes were employed. RsfS successfully dissociated non-crosslinked 70S ribosomes into subunits and formed a stable complex with 50S subunits, as detected by immunoblot analysis of 50S subunits after sucrose gradient centrifugation (Fig. S2a). In contrast, RsfS was unable to dissociate crosslinked 70S ribosomes and consequently did not inhibit oligo-Phe synthesis (Fig. S2b). If RsfS were capable of dissociating any 70S ribosomes, the cell would die due to the inability to synthesize proteins. This would make RsfS a highly toxic factor, contradicting its conservation and essential role in cell survival during starvation.

Next, we addressed the question: does RsfS dissociate any of the 70S ribosomes, or does it have a specific target? Vacant 70S ribosomes were reconstituted by incubating 50S and 30S subunits in association buffer, and then incubated either in the absence or presence of RsfS. Sucrose gradient ultracentrifugation showed that RsfS efficiently dissociated these empty 70S ribosomes into their subunits (Fig. 4c). However, when the ribosomes were manipulated into Pi, PRE, and POST states, RsfS was no longer able to dissociate any of these ribosomes (Fig. 4d). Whenever the 70S ribosomes were occupied by tRNAs (translating 70S ribosomes), RsfS was unable to dissociate them, thus resulting in no inhibition of translation in a Poly(U)-dependent oligo (Phe) synthesis assay where most of the 70S ribosomes were engaged in translation (Fig. 4e). However, when the 50S subunits were pre-incubated with RsfS, protein synthesis was blocked (Fig. 4f) due to its inhibition of subunit association. Overall, RsfS exclusively dissociates idle 70S ribosomes and subsequently prevents the reassociation of ribosomal subunits, without affecting actively translating ones, thereby blocking translation activity from both canonical and 70S form initiation (Orelle et al., 2015; Yamamoto et al., 2016).

### RsfS inhibits EF-G dependent GTPase activity triggered by vacant 70S ribosomes

The energy deprivation observed in the *ΔrsfS* strain could be attributed to the incapacity of these cells to dissociate vacant 70S ribosomes, resulting in a slower adaptation to starvation conditions. To investigate whether vacant 70S ribosomes consume energy, the EF-G dependent GTPase activity was measured in the presence of either vacant 70S ribosomes or ribosomal subunits. It was observed that EF-G could not hydrolyze GTP in the presence of either 30S or 50S subunits alone (Fig. S3). However, in the presence of both subunits, EF-G dependent hydrolysis of GTP occurred at a rate of 74.5 GTPs per ribosome per minute. This level of GTPase activity is comparable to that observed when vacant 70S ribosomes were present (83.6 GTPs per ribosome per minute). With increasing concentrations of RsfS, the GTPase activity of EF-G decreased in both 70S ribosomes and autonomously assembled subunits in a dose-dependent manner (Fig. 5). These findings indicate that RsfS dissociates vacant 70S ribosomes and inhibits autonomous subunit assembly, leading to a reduction in EF-G GTPase activity and thereby conserving energy.

**Figure 5:**
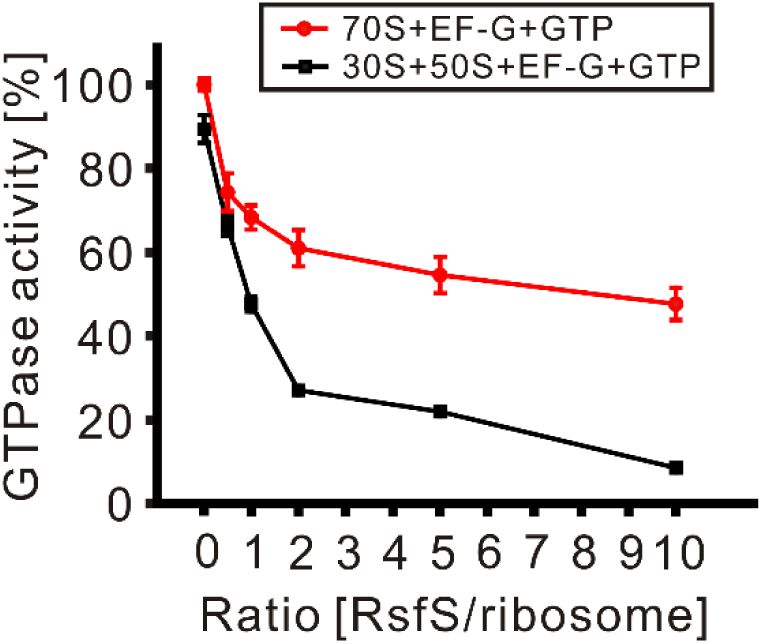
RsfS inhibits EF-G dependent GTPase activity. EF-G dependent GTPase activity of 70S and (30S+50S) respectively was determined in binding buffer, *via* malachite green assay. RsfS was titrated in 0-10 folds excess over ribosomes in molar ratio. Data are presented as mean values +/- S.D. based on n=2 technical replicates. For details see Materials and methods.

## Discussion

During nutrient downshift, bacteria must activate synthetic amino acid pathways to produce essential amino acids that they can no longer acquire from the environment. Additionally, non-essential protein synthesis and metabolic activities must be downregulated to conserve cellular resources and energy. (Andersson & Kurland, 1990; Dong, Nilsson, & Kurland, 1996). One well studied mechanism that bacteria use to adapt to these conditions is the stringent response. In *E. coli*, this response has been shown to act at the level of transcription; the alarmones (p) ppGpp produced by RelA directly inhibit transcription from ribosomal promoters, resulting in reduced expression of ribosomal proteins. Meanwhile, the alarmones increase the expression of genes involved in amino acid biosynthesis (review see ref. (Potrykus & Cashel, 2008)). Although the stringent response inhibits *de novo* ribosome synthesis, it does not regulate pre-existing ribosomes. It has been reported that reduction of the number of translating ribosomes enables *E. coli* to maintain the elongation rate during slow growth in nutrient poor medium (Dai et al., 2016). Here, we demonstrated the detailed mechanism that RsfS plays a prominent role in this regulation of nutrient deprivation by silencing ribosome activities. Strains lacking of *rsfS* leads to a severe delay of adaptation during the nutrient downshift, exhibiting its physiological importance. Our results highlighted multifaceted functions of RsfS to save cellular energy and resources (Fig. 6):

1. RsfS reduces GTPase activity trigged by ribosomes. After nutrient downshift, in strains lacking *rsfS*, there was a substantial accumulation of idle 70S ribosomes *in vivo* (Fig. 2d), which resulted in a waste of energy, as EF-G continues to be able to bind and hydrolyze GTP, at a rate of 74 ± 23 s^-1^ per ribosome (Rodnina, Savelsbergh, Katunin, & Wintermeyer, 1997). This GTPase activity is even higher than that observed when bacteria are grown in nutrition rich medium. When the cells are in the log-phase, two GTPs are hydrolyzed per elongation cycle: one by EF-G and one by EF-Tu (Cabrer, San-Millian, Vazquez, & Modolell, 1976). When almost all the ribosomes present in the cell are engaged in protein synthesis, as is the case when in log-phase, EF-G hydrolysis of GTP occurs at the rate of the elongation cycle, namely 10 to 20 s^-1^ (H. Bremer & Dennis, 1996). RsfS blocks the EF-G dependent GTPase activity with vacant 70S ribosomes and the inhibition is even stronger with 30S and 50S subunits (Fig. 5), to save the cellular energy in the poor medium (Fig. 3b).
2. RsfS reduces the translation activity. Our result showed that RsfS reduces the translation activity in cell only if vacant 70S are dissociated and the 50S subunits can bind RsfS before they enter the process of proteins synthesis, but protein synthesis is not perturbed by RsfS when 70S ribosomes that already contain tRNA (Fig. 4d). This is evidenced by the fact that RsfS does not affect the viability of log-phase *E. coli* cells (Häuser et al., 2012). Therefore, the constant presence of RsfS is an “always-ready” backup for possible nutrition shortage.

**Figure 6:**
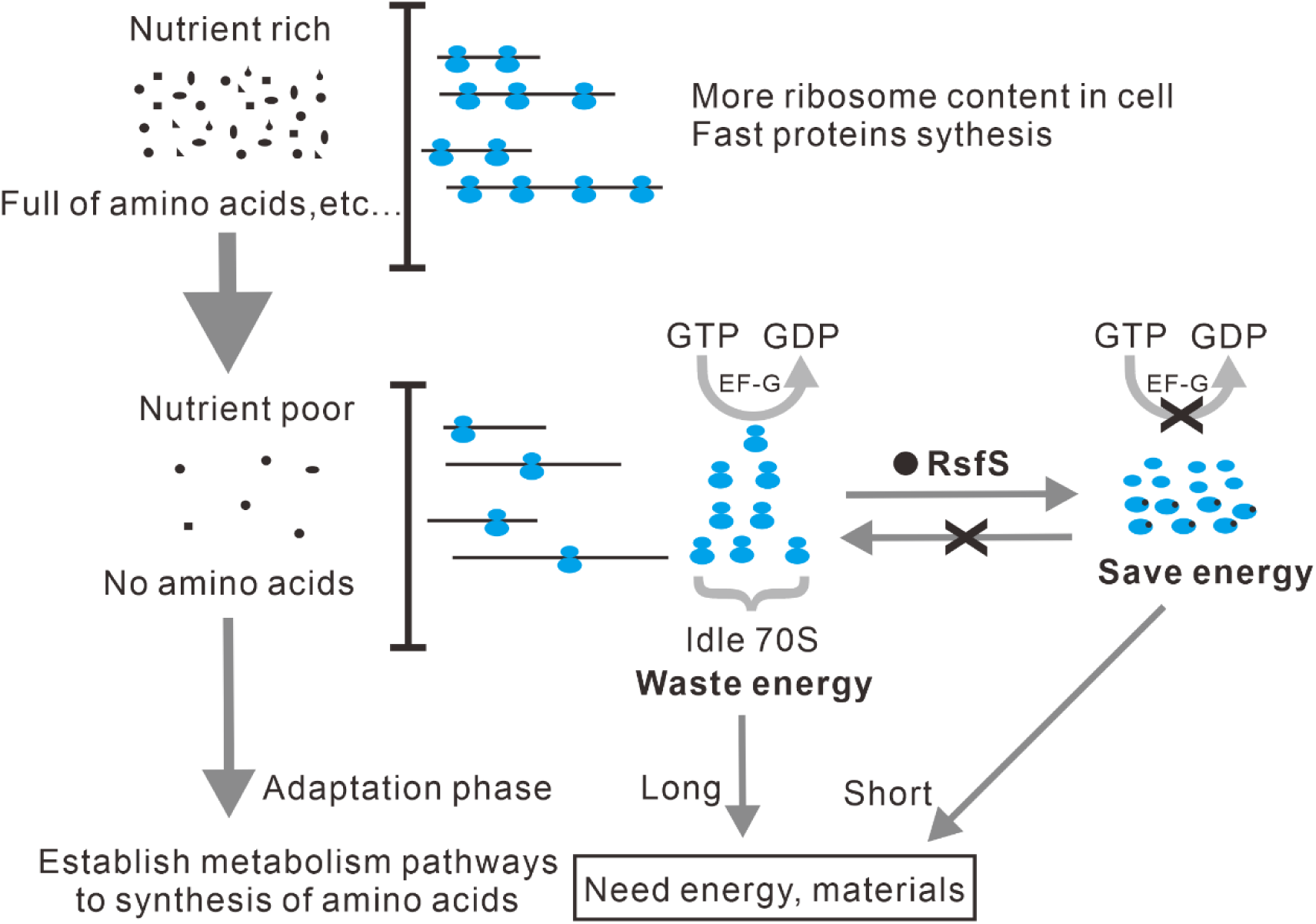
Model for the contribution of RsfS to metabolic adaptation upon nutrient deprivation in bacteria. Under nutrient rich condition (upper case), bacterial cells take up directly many metabolic building blocks in particular peptides from the medium, thus the cells have high ribosome content to produce proteins at maximum rates. However, in nutrient poor condition (lower case), many synthetic pathways involved in the production of amino acids have to be established, which costs cellular resource and energy. But at the same time, the accumulation of idle 70S ribosomes cause a devastating energy drain by stimulating the activity of GTPase proteins such as EF-G. RsfS dissociates the idle 70S ribosomes and prevents the reassociation of ribosome subunits into 70S ribosomes to save energy and cellular materials, so that bacteria can adapt nutrient poor condition fast. Lack of RsfS results in a severe delay in adapting the nutrient poor condition.

What is the intrinsic reason for RsfS exclusively dissociating vacant 70S ribosomes while sparing those involved in programmed translation (Pi, PRE, and POST states)? We have identified steric clashes between RsfS (in the 50S-RsfS complex) and its binding site on translating 70S ribosomes, which prevent the binding of RsfS to these ribosomes. Interestingly, similar steric hindrances were also observed between RsfS and vacant 70S ribosomes. Moreover, the binding site of RsfS does not overlap with any translation factors, thus ruling out a competition mechanism. One plausible explanation could be that vacant 70S ribosomes (lacking tRNAs) exhibit a lower binding affinity of subunits compared to translating ones, potentially facilitating RsfS’s access to its binding site at the 50S subunit interface and subsequent dissociation of the vacant 70S ribosomes.

There could be other explanations for the inability of *ΔrsfS* strain adapts to the nutrient downshift. Bacterial RsfS was initially identified as a protein migrating close to 50S subunits in a sucrose gradient, using a semi-quantitative mass spectrometry approach, suggesting a possible role in 50S assembly (Jiang et al., 2007). In addition, the human mitochondrial homolog MALSU1 was found in a cryo-EM based structural approach analyzing mitochondrial large ribosomal subunit precursors and was suggested to be an assembly factor preventing participation of immature large subunits in translation (Brown et al., 2017). The initiation factor eIF6 also docks to the ribosomal protein uL14 of eukaryotic 60S ribosomes like RsfS. It was reported as a 60S assembly factor and plays an essential role in the late pre-25S rRNA processing and the export of the 60S subunit from the nucleolus to the cytoplasm (Biswas et al., 2011). It seems that eIF6 and RsfS are involved in different aspects of the ribosomal life cycle. Hence, it is attractive to speculate that both eIF6 and RsfS are involved in the allocation of large ribosomal subunits to achieve a partitioning of ribosomal activity that matches the cellular demand.

This study illustrated how RsfS enables cells to efficiently utilize cellular resources, complementing the stringent response, thereby enabling bacteria to adapt to nutrient deprivation conditions. It is also possible that the inability of the *ΔrsfS* strain to trigger the stringent response contributes to the observed growth problem. i.e. RsfS works in line with the stringent response and thus plays a key role in the translational regulation during the nutrient deprivation. The broader implications of those results are on research conducted on ribosome shutdown mechanisms which accrue in human pathogen infection processes, where the host restrains pathogen from proliferating by imposing various of stresses, such as *S.aureus* and *B. burgdorferi* (Khusainov et al., 2020; Prusa, Zhu, & Stallings, 2018; Sharma et al., 2023). Therefore, bacterial RsfS can serve as a potential drug target for antimicrobial treatments.

## Materials and methods

### Strains

*E. coli* BW25113 was used as wild type (WT) strain, and the *rsfS* deletion strain JW5090-2 was obtained from Keio collection (Baba et al., 2006). We also made a rescue strain based on the *ΔrsfS* strain, in which RsfS was overexpressed (Häuser et al., 2012).

### Overproduction and purification of recombinant His_6_RsfS

Our first preparations of RsfS using standard procedures for His-tagged recombinant proteins expressed in *E. coli* were contaminated with GTPase activities. The following procedure result in highly purified RsfS proteins completely free of GTPase contaminants.

Chemical competent BL21 (DE3) pLysS cells were transformed with pHGWA-*ybeB* plasmid (Häuser et al., 2012) and grown in LB medium containing 50 µg/ml ampicillin (Amp) for 15 hours at 37°C. Cells were sedimented, washed with LB medium and a 2xYT culture supplemented with Amp was inoculated to a start OD_600_ of 0.13. Cells were grown in baffled flasks at 200 rpm and 30°C for 3 hours. At OD_600_ of 0.8 production of recombinant His_6_RsfS was induced with 1 mM IPTG. After 3 hours of incubation at 200 rpm and 30°C cells were sedimented by centrifugation (in a JLA9.100 rotor for 20 min at 5,000 rpm and 4°C), resuspended in lysis buffer (50 mM TRIS pH 8, 200 mM NaCl, 10% glycerol, 10 mM Imidazole (IZ), TM complete EDTA-free protease inhibitor (Roche)), frozen in liquid N_2_ and stored at −80°C. Cells were thawed on ice and lysed by three microfluidizer cycles. Cell lysates were cleared by centrifugation (30 min at 45,000 rpm at 4°C). Ni-NTA slurry (Qiagen) (0.8 ml bed volume) was equilibrated with lysis buffer, mixed with cleared cell lysate and incubated for 2 hours on a rotating wheel at 4°C. The mixture was poured into a PD10 column (GE) and flow through was collected by gravity flow. Beads were washed with 20 ml wash buffer (50 mM TRIS pH 8.0, 500 mM NaCl, 0.2% Triton X-100, 30 mM IZ, 10% glycerol, TM complete EDTA free), 10 ml buffer A (50 mM TRIS pH 8.0, 300 mM NaCl, 10 mM IZ, 10% glycerol), 10 ml buffer B (buffer A + 1% Triton X-100), 10 ml buffer C (buffer A + 1 M NaCl), 10 ml buffer D (buffer A + 10 mM MgCl_2_ + 3 mM ATP), 10 ml buffer E (buffer A + 0.5 M TRIS pH 8.0) and 10 ml buffer A with 10% elution buffer. 250 µl elution buffer (50 mM TRIS pH 8.0, 300 mM NaCl, 250 mM IZ, 10% glycerol, TM complete EDTA free) were added and immediately collected (void volume). Protein was eluted by addition of 1 ml elution buffer after 20 min incubation on a rotating wheel (E1), followed by a second 1 ml elution step (E2). Samples of the eluted fractions where analyzed by a 15% SDS polyacrylamide gel electrophoresis (PAGE) stained with Coomassie solution E1 and E2 were pooled and loaded on a Superdex 75 High load 16/60 column (GE) equilibrated with HMK buffer (20 mM HEPES pH 7.6, 6 mM Mg(Ac)_2_, 150 mM KAc, 10% glycerol, 4 mM β-mercaptoethanol). The column was run with a flow rate of 1 ml/min and fractionation was started after 40 ml. His_6_RsfS eluted in a major peak at app. 75 ml. Fractions covering the elution peak were analyzed by SDS-PAGE. Fractions of highest concentration and purified to apparent homogeneity were aliquoted, flash frozen in liquid N_2_ and stored at −80°C.

### Growth and viability test of WT and ΔrsfS strain after a nutrient shift-down

WT and *ΔrsfS* cells were grown overnight in LB medium and then diluted 1:1000 in M9 minimal medium (1 liter contains 12.8 g Na_2_HPO_4_•7H_2_O, 3 g KH_2_PO_4_, 0.5 g NaCl, 1 g NH_4_Cl, 0.4% glucose, 2 mM MgSO_4_, 0.1 mM CaCl_2_) to a start OD_600_ of 0.005. Two cell suspensions (duplicates derived from an overnight culture) of 25 ml were grown in 100 ml flasks at 37°C and a shaking frequency of 200 rpm for 42 hours. Samples were taken at indicated points of time. Cell density was determined photometrically (OD_600_) and after appropriate dilution cell suspensions were spread on LB agar plates. The plates were incubated at 37°C for 12 hours. Single colonies were counted and initial concentrations of viable cells were calculated. The two growth curves appeared to be almost identical, hence we used one as a representative example.

### 70S ribosome loading analysis

A ribosome associated with mRNA would produce a ribosome-protected fragment (RFP) in Ribo-seq. To normalize the initial material, PCR cycles and the fraction of RFP reads among all sequencing reads, we loaded 10 µl of Ribo-seq library for each sample on an 8% polyacrylamide gel with 3 µl marker. The intensities of bands were quantified with background subtracted. More PCR cycles would lead to higher intensity. Higher rate of the Ribo-seq reads that are mapped to the mRNAs reflects more ribosome occupancy. Therefore, the 70S ribosome loading was estimated as:

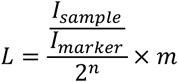

Where I is the intensity (for samples, only the RFP library band of 147-160bp were measured), n is the PCR cycle number, and m% is the mapping rate to the mRNAs.

### Differential gene expression analysis

Differentially expressed genes were analyzed using edgeR package (Robinson, McCarthy, & Smyth, 2009). We considered genes with |log_2_ fold change|>1 and p<0.05 as differentially expressed genes. We used PANTHER v13.0 (http://pantherdb.org/) to conduct gene ontology (GO) overrepresentation analysis of up-/down-regulated genes, with the significance threshold of FDR<0.01. The modules of the overrepresented genes were visualized using Ecocyc database (https://ecocyc.org/).

### Ribosome profiling

*E. coli* strains were cultivated in M9 minimal medium at 37°C in flasks. Chloramphenicol was added 15 min before harvest to the bacterial culture to the concentration of 100 µg/mL. The cells were harvested by centrifugation at 5000 x g for 10 min at 4°C, and then were washed with pre-chilled PBS in presence of 100 µg /mL Chloramphenicol. Cell pellet was then gently resuspended in 6 mL pre-chilled sucrose buffer (16 mM Tris buffer pH=8.1, supplemented with 0.5 M RNase-free sucrose, 50mM KCl, 8.75 mM EDTA, 100 µg/mL chloramphenicol and 12.5 mg/mL lysozyme). Then the cells were again pelleted by centrifugation at 5000 x g, 10 min. Up to 0.6 mL of cell lysate was layered onto 35% (w/w) sucrose buffer (10 mM Tris buffer pH=7.8 containing 50 mM NH_4_Cl, 10 mM MgCl_2_, 0.2% Triton X-100 and 100 µg/mL chloramphenicol) and ultracentrifuged at 42500 rpm for 5 h in type 70 Ti rotor (Beckman Coulter) at 4°C. The pellet was gently resuspended in 0.2 mL micrococcal buffer (50 mM Tris buffer pH=7.9 containing 5 mM CaCl_2_ and 12 mM MgCl_2_). An aliquot of 1 A_260_ unit of ribosome-bound mRNA fraction was digested with 800 units Micrococcal nuclease (NEB) for 20 min at 25°C. RNA was extracted using Trizol method (Rio, Ares, Hannon, & Nilsen, 2010). Ribo-seq libraries were prepared using standard VAHTS^TM^ Small RNA Library Prep Kit for Illumina (Vazyme). Sequencing was performed on an Illumina HiSeq X Ten sequencer. Raw datasets have been deposited to GEO database (accession number GSE142817).

Adapter sequences were trimmed from the high-quality reads. The clean reads were mapped to E. coli reference genome sequences (NCBI accession number NC_000913.3) using FANSe3 algorithm (Liu, Xiang, Zheng, Jin, & Zhang, 2018) with parameters -E2 --indel -S10 -U1. Genes with at least 10 mapped reads were considered quantifiable (Bloom, Khan, Kruglyak, Singh, & Caudy, 2009). Gene expression levels were quantified using RPKM method (Mortazavi, Williams, McCue, Schaeffer, & Wold, 2008).

### Cell lysate based rapid translation system (RTS)

The rapid translation system has been performed according to the producer’s protocol with slight modifications (Roche, Rapid Translation System RTS 100, *E. coli* HY Kit). Standard volume of reaction batch containing reconstitution buffer, reaction mix, amino acids and DNA has been increased from 10 µl to 20 µl. Adaptation buffer was used for adjusting the reaction mixture to binding buffer (20 mM Hepes-KOH pH 7.6 (0°C), 4.5 mM Mg (Ac)_2_, 150 mM KAc, 4 mM β-mercaptoethanol, 2 mM spermidine and 0.05 mM spermine), which was optimized for efficient protein synthesis (Szaflarski et al., 2008). As a reporter, a DNA plasmid containing a Renilla luciferase gene was used. The reaction was stopped and quenched after 1h of incubation at 30°C. Chemoluminescence was measured for 30 seconds.

### Dissociation test of empty 70S or 70S programmed with tRNA (Pi, PRE and POST states)

For empty 70S, equal molar ratio of 30S and 50S were incubated for 20 min at 40°C in association buffer (20 mM Hepes-KOH pH 7.6 (0°C), 20 mM Mg(Ac)_2_, 30 mM KAc, 4 mM β-mercaptoethanol), which allows the ribosomal subunits associate into 70S, and then the re-associated 70S was isolated by sucrose-gradient centrifugation. 5 pmol of this newly prepared empty 70S and 50 pmol RsfS were incubated for 30 min at 37°C in binding buffer.

For programmed 70S, the pre-incubation reaction containing 5 pmol of 70S programmed with 40 pmol of MF mRNA (Triana-Alonso, Dabrowski, Wadzack, & Nierhaus, 1995) and 10 pmol of deacylated-tRNA^fMet^ was carried out in binding buffer for 15 min at 37°C (pi-translocational (Pi) state). Then 10 pmol of N-acetyl-[^14^C]Phe-tRNA^Phe^ ([^14^C]-Ac-Phe-tRNA^Phe^),and – if indicated - 50 pmol RsfS was added to the reaction and incubated for 30 min at 37°C (pre-translocational (PRE) state). Subsequently GTP (2.5 mM f.c.) and 4 pmol of EF-G was added and translocation of A- and P-tRNA to P and E sites was performed for 5 min at 37°C (post-translocational (POST) state) (Hausner, Geigenmüller, & Nierhaus, 1988). The efficiency of translocation was measured by the puromycin reaction and the tRNA occupancy was calculated using nitrocellulose filtration (Blaha et al., 2000).

After each incubation step two aliquots were withdrawn from the main reaction and subjected to a sucrose-gradient centrifugation. The ribosome profile was analyzed at 260 nm.

### Poly(U)-dependent oligo (Phe) synthesis with precharged Phe-tRNA

70S programmed was obtained by incubation of 3 pmol empty 70S with 30 µg poly(U) mRNA and 6 pmol acylated-tRNA^Phe^ for 10 min at 37°C, prior to the incubation with or without 30 pmol RsfS for 10 min at 37°C in 15 µl binding buffer. 3 pmol of 50S subunits were incubated with 30 µg poly(U) mRNA with or without 30 pmol RsfS for 10 min at 37°C in 15 µl binding buffer. The sample was further incubated with 3 pmol 30S subunits for 10 min at 37°C and then analyzed in poly(U)-dependent oligo (Phe) synthesis (Szaflarski et al., 2008).

For poly(U)-dependent oligo (Phe) synthesis, 2.4 pmol of EF-G together with the ternary complex mix were added yielding 30 µl in binding buffer. The ternary complex mix contained in 15 µl 30 pmol [^14^C]Phe-tRNA^Phe^, 45 pmol EF-Tu, 45 pmol EF-Ts, 3 mM GTP and was preincubated 5 min at 37°C. Afterwards samples were incubated at 30°C for 10 min and 10 µl aliquots were precipitated with TCA, incubated at 90°C for 5 min in the presence of 30 µl of 1% (w/v) BSA and filtered through glass filters. Activity of [^14^C]Phe-tRNA^Phe^ was counted using a scintillation counter.

### Sucrose gradient centrifugations

Centrifugation of 70S loading test was performed in SW40 rotor for 19 h at 21,000 rpm, 10-40% sucrose gradient in Tico buffer (20 mM Hepes-KOH pH 7.6 (0°C), 6 mM Mg (Ac)_2_, 30 mM K(Ac), 4 mM β-mercaptoethanol). For RTS and oligo (Phe) synthesis experiments, centrifugation was performed in SW60 rotor for 2 h at 38,000 rpm, 10-30% sucrose gradient in binding buffer. Centrifugation of 70S dissociation test was performed in SW40 rotor for 20 h at 22,000 rpm, 10-30% sucrose gradient in binding buffer. A_260_ was measured to detect ribosomal RNA during pumping out.

### EF-G dependent GTPase activity in the presence of 70S and (30S+50S) subunits

Malachite Green Reagent: 50 mg malachite green (final concentration 1 mg/ml) and 500 mg ammonium molybdate tetrahydrate (final concentration 10 mg/ml) were dissolved in 50 ml of 1 M hydrochloric acid (HCl) followed by filtration through a 0.45 nm filter. The solution was stored at room temperature in the dark for a maximum of 2 weeks.

GTPase assay: Indicated amounts of RsfS were incubated with ribosomes (3 pmol of each) in binding buffer at 37°C for 10 min. GTP (final concentration 0.2 mM) and EF-G (6 pmol) were added to the reaction, followed by an incubation at 37°C for 5 min. 35 µl aliquot of the reaction above was mixed with 75 µl of the Malachite Green solution for 10 min at room temperature and the absorbance at 650 nm was monitored. A standard curve from 10 to 400 µM Pi (KH_2_PO_4_) was generated for each experiment and read in parallel. Samples without ribosomes were used as blank control, the corresponding value was subtracted. The data were processed with the software Origin8 (OriginLab Corporation).

## Acknowledgements

Firstly, we thank late Prof. Knud Nierhaus with our deep respect for his supervision, Institut für Medizinische Physik und Biophysik (IMPB), Charité–Universitätsmedizin Berlin. Thanks also go to Prof. Christian Spahn (IMPB), Dr. Matthew Kraushar, Max-Planck-Institut für Molekulare Genetik (MPIMG), Berlin for intensive scientific discussions, excellent suggestions, Renate Albrecht (MPIMG) for ribosome preparations. This work was supported by the National Key Research and Development Program of China [2023YFA0915800], Guangdong Key Research and Development Program [2019B020226001] and the Max-Planck-Institut für Molekulare Genetik, Berlin, for financial support.

## Author contributions

B.Q. and G.Z. conceived the project; B.Q., H.Z., R.N., L.J. and Y.B. performed the experiments; B.Q., H.Z., R.N. and G.Z. analyzed the data; B.Q. and G.Z. wrote the paper.

## Competing Interests

The authors declare no competing interests.

## Notes

### Competing Interest Statement

The authors have declared no competing interest.

### Summary of Updates

In this revised version, we have updated our manuscript to incorporate recent advances in the field of ribosome hibernation and RsfS function. Since the initial posting of our preprint in 2024, several key studies have emerged that contextualize and strengthen our findings. Helena-Bueno et al. (2024) provided a comprehensive overview of bacterial ribosome hibernation factors, highlighting that most known factors primarily stabilize inactive ribosomes, whereas mechanisms for selectively eliminating vacant 70S ribosomes remain largely unexplored. This further underscores the significance of our discovery that RsfS serves as the sole factor capable of selectively dissociating vacant 70S ribosomes. More recently, Schmid et al. (2026) demonstrated that RsfS (Lpg1377) functions as a ribosome assembly inhibition factor in the intracellular pathogen Legionella pneumophila, coordinating with other hibernation factors to support starvation survival, host cell infection, and antibiotic tolerance. This study provides independent physiological evidence for RsfS-mediated ribosome regulation in a pathogenic context, complementing our mechanistic findings in Escherichia coli.

